# Conditional interactions in literature-curated protein interaction databases

**DOI:** 10.1101/352328

**Authors:** R. Greg Stacey, Michael A. Skinnider, Jenny H. L. Chik, Leonard J. Foster

## Abstract

Databases of literature-curated protein-protein interactions (PPIs) are often used to interpret high-throughput interactome mapping studies and estimate error rates. These databases combine interactions across thousands of published studies and experimental techniques. Because the tendency for two proteins to interact depends on the local conditions, this heterogeneity of conditions means that only a subset of database PPIs are interacting during any given experiment. A typical use of these databases as gold standards in interactome mapping projects, however, assumes that PPIs included in the database are indeed interacting under the experimental conditions of the study. Using raw data from 20 co-fractionation experiments and six published interactomes, we demonstrate that this assumption is often false, with up to 55% of purported gold standard interactions showing no evidence of interaction, on average. We identify a subset of CORUM database complexes that do show consistent evidence of interaction in co-fractionation studies, and we use this subset as gold standards to dramatically improve interactome mapping as judged by the number of predicted interactions at a given error rate. We recommend using this CORUM subset as the gold standard set in future co-fractionation studies. More generally, we recommend using the subset of literature-curated PPIs that are specific to experimental conditions whenever possible.

## Introduction

Proteins perform the majority of cellular functions necessary for life. Nearly all individual proteins are modular components of larger macromolecular structures, i.e. protein complexes, and the exact role of a protein within a cell is controlled by its interactions with co-complex members. Uncovering which proteins interact, i.e. the interactome, is therefore central to understanding the molecular mechanisms of life.

This task is complicated by a combinatorial explosion, however: a proteome containing 20000 proteins has nearly 200 million potential pairwise interactions and many more higher order complexes. High-throughput techniques that analyze thousands of proteins simultaneously with minimal bias offer a solution to this problem (1). For example, PCP-SILAC (protein correlation profiling-stable isotope labeling of amino acids in cell culture), a co-fractionation (CF) technique, separates protein complexes into fractions according to their size (rotational cross-section), and associates proteins whose amounts are correlated between fractions. As each fraction is analyzed with mass spectrometry, PCP-SILAC and other CF techniques can detect tens of thousands of interacting proteins (2–8). In order to separate signal from noise, it is common for high-throughput protein interactome studies to consult databases of known, unequivocal interactions (“gold standards”) (2,3,9,10). For example, co-fractionation studies often use gold standard interactions as training labels in a machine learning classifier (2,3,11). Gold standards are also used to define false positive/negative and true positive/negative interactions in order to calculate common statistics such as precision, recall, and sensitivity (5–7,11,12).

Gold standard databases are assembled from different experiments and techniques, each with a unique set of biases. Since protein-protein interactions (PPIs) can be conditional and transient, single datasets, which are typically generated by a single technique, can disagree with gold standards. This variability partly reflects true biological differences. For example, the majority of *in vivo* yeast PPIs were observed to depend on environmental and chemical conditions (13). Some assays also impose technical biases that limit detectable PPIs, such as a bias of some high-throughput techniques toward highly expressed or well studied protein pairs, or a bias against PPIs involving transmembrane proteins (12). Therefore, gold standard databases that include all interactions that can occur will fail to describe the subset of interactions that are either not occurring due to current experimental conditions, or that are unlikely to be detected due to technical limitations.

Therefore, a distinction should be made between the large, curated compilations of interactions across many studies, and the gold standard sets used as a reference for a single dataset. Our own focus has been on interactome mapping using co-fractionation, so here we quantify the proportion of gold standard interactions that fail to display any evidence for interaction in 20 co-fractionation datasets. Using a conservative measure of protein interaction, we find that between 19 and 55% of gold standard PPIs display no evidence of interacting. Across co-fractionation experiments, there is evidence that a subset of literature-curated complexes consistently co-fractionates, suggesting this subset would be a more appropriate gold standard reference set. Indeed, the number of predicted interactions at a given stringency increases dramatically when using this subset as a gold standard set. We recommend using this subset as the gold standard reference in future co-fractionation studies and, more generally, using experiment- and condition-specific gold standards whenever possible.

## Results

### 2.1 Discrepancies exist between gold standards and individual datasets

Using the CORUM database of protein complexes (14), we first examined the degree to which literature-curated PPIs were unsupported by data from single co-fractionation datasets. Many database PPIs show clear evidence of interaction, as shown by their tendency to co-fractionate for the entire chromatogram (Fig 1A) or a portion of the chromatogram (Fig 1B). However, other protein pairs from within a single CORUM complex show little evidence of interaction in certain experiments. For example, two chaperone proteins, HSP-90a (UniProt ID P07900) and BiP (P11021) are known to interact as part of a larger chaperone multiprotein complex (15) (CORUM complex “HCF-1”), yet there is little evidence that the two proteins co-fractionate in our data (Fig 1C).

**Fig 1.**
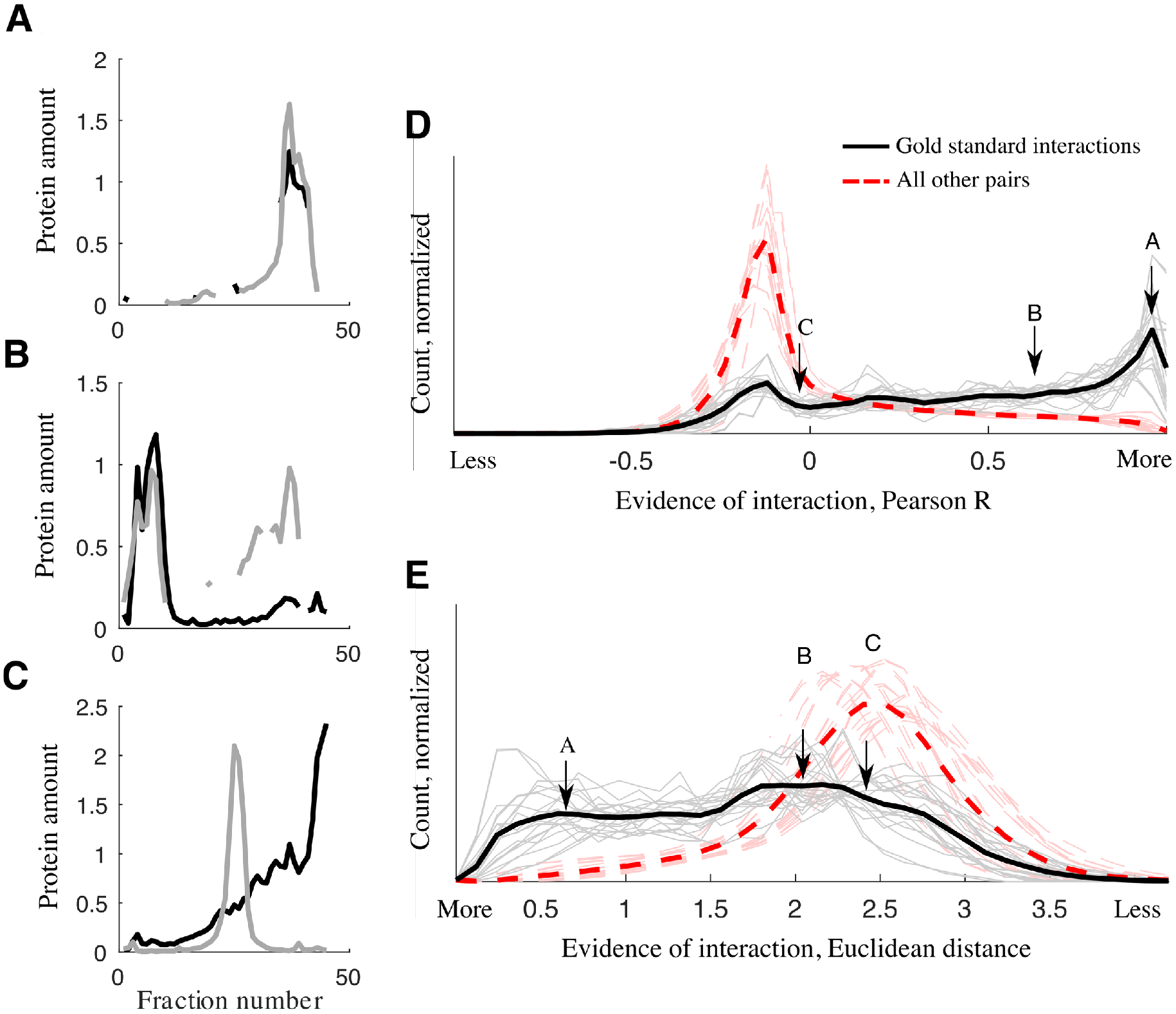
Not all CORUM gold standard interactions are supported by co-fractionation data. A.
Example gold standard pair with strong evidence for interaction. Q9NQP4 and E5RGS4, prefoldin complex. B. Example gold standard pair with evidence for interaction. Q14103 and O75534, PIN1-AUF1 complex. C. Example gold standard protein pair with little data-derived evidence for interaction. P11021 and P07900, HCF-1 complex. D. Histogram of Pearson correlation coefficients and E. Euclidean distance between every gold standard interaction in our co-fractionation data (20 datasets, grey; average, black). All other protein pairs in our data are shown, the vast majority of which are not interacting (red). Example pairs A, B, C are shown (arrows).

More broadly, across 20 PCP-SILAC co-fractionation datasets, the majority of random protein pairs do not co-fractionate, as quantified by anti-correlated fractionation profiles, a conservative measure of which protein pairs are non-interacting (red, Fig 1D). While the majority of gold standard protein pairs have positively correlated co-fractionation profiles (black), 23% (34442/149477) are negatively correlated. All 20 datasets include a similar proportion of negatively correlated gold standard pairs (23 +/− 5%, mean +/− st.d.). This pattern is similar when co-fractionation is measured with Euclidean distance, another standard measure (Fig 1E).

While the full set of CORUM complexes is a widely used gold standard (6,7,9–11), there are many other literature-curated interaction databases. In addition to CORUM, we examined nine databases of protein interactions (16–24) and two subsets of CORUM used previously as gold standards (2,3). These range from databases that include interactions from high-throughput experiments to manually curated databases composed exclusively of low-throughput experiments. All had anti-correlated protein pairs in our co-fractionation datasets (Fig 2). As a baseline, 62% of all protein pairs, the large majority of which can be assumed to be non-interacting, were anti-correlated (Fig 2, red). The proportion of anti-correlated pairs in gold standard sets ranged from 55% (HPRD) to 19% (CORUM). Restricting gold standard PPIs to those supported by two or more publications limits but does not eliminate uncorrelated protein pairs (Supp. Fig 1). Therefore all interaction databases investigated here contain protein pairs that are not supported by our co-fractionation data, and comparisons to CORUM give a conservative estimate of the discrepancy between our data and interaction databases.

**Fig 2.**
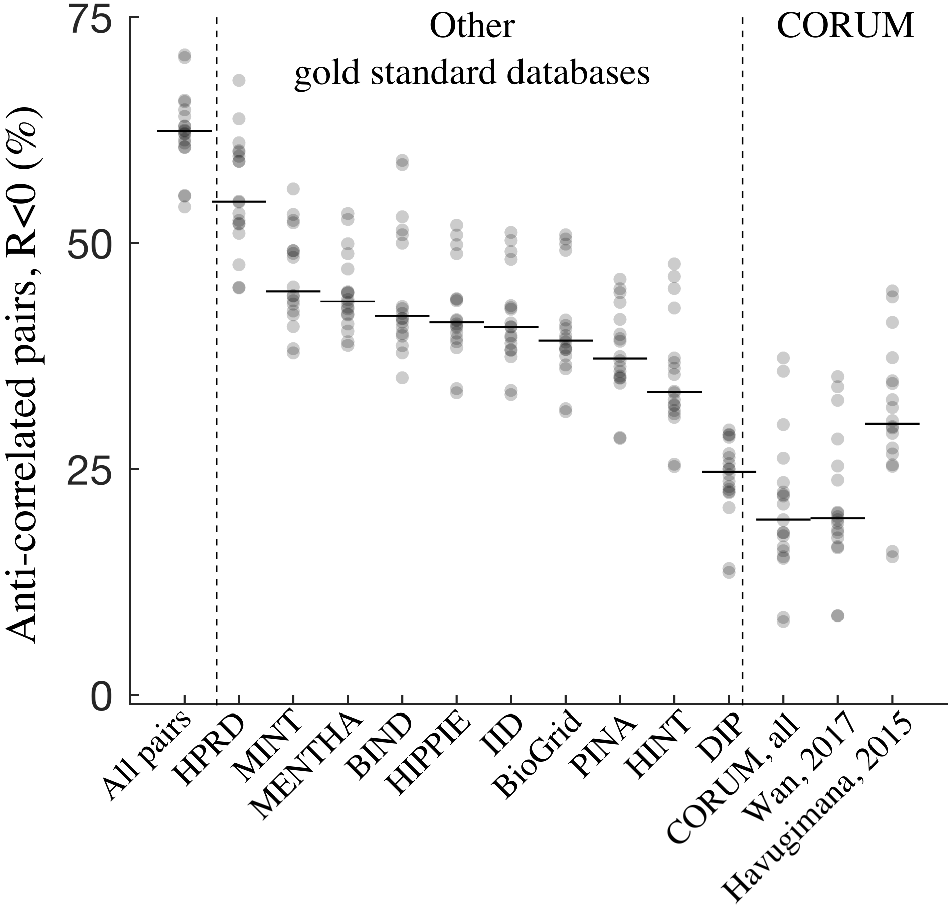
Protein pairs across many gold standard databases do not co-fractionate, as measured by anti-correlated co-fractionation profiles (Pearson correlation R<0). Each point is one dataset. Horizontal lines show medians. Red: all non-gold standard protein pairs. Only non-redundant gold standard pairs were analyzed.

### 2.2 Discrepancies are consistent within and between high-throughput techniques

A certain level of discrepancy between raw co-fractionation profiles and interaction databases should be expected, as all experimental samples undergo some degradation owing to the constraints imposed by each assay. But do individual datasets differ randomly or systematically from interaction databases? For certain gold standard complexes we see a strong tendency for co-complex members to co-fractionate, and, conversely, a strong tendency for other complexes to fail to co-fractionate (Fig 3A). Across 20 co-fractionation datasets, we made 39846 pairwise comparisons between fractionation profiles of cytoplasmic ribosomal proteins, and 17616 pairwise comparisons of proteins in the C complex spliceosome. The collection of ribosomal gold standard interactions are significantly better correlated than chance (R = 0.64, chance R = 0.48, p = 0.005, permutation test; Fig 3B), while the collection of spliceosome gold standard interactions are significantly worse correlated (R = 0.27, p < 0.001; Fig 3C). Calculating significance for all 1253 observed CORUM complexes (permutation test, Benjamini-Hochberg correction), 419/1253 correlate significantly higher than average, while 294/1253 are significantly lower. This suggests that some gold standard complexes are enriched for interacting protein pairs, while others are enriched for non-interacting protein pairs, where non-interacting pairs likely represent interactions disrupted by the particular assay.

**Fig 3.**
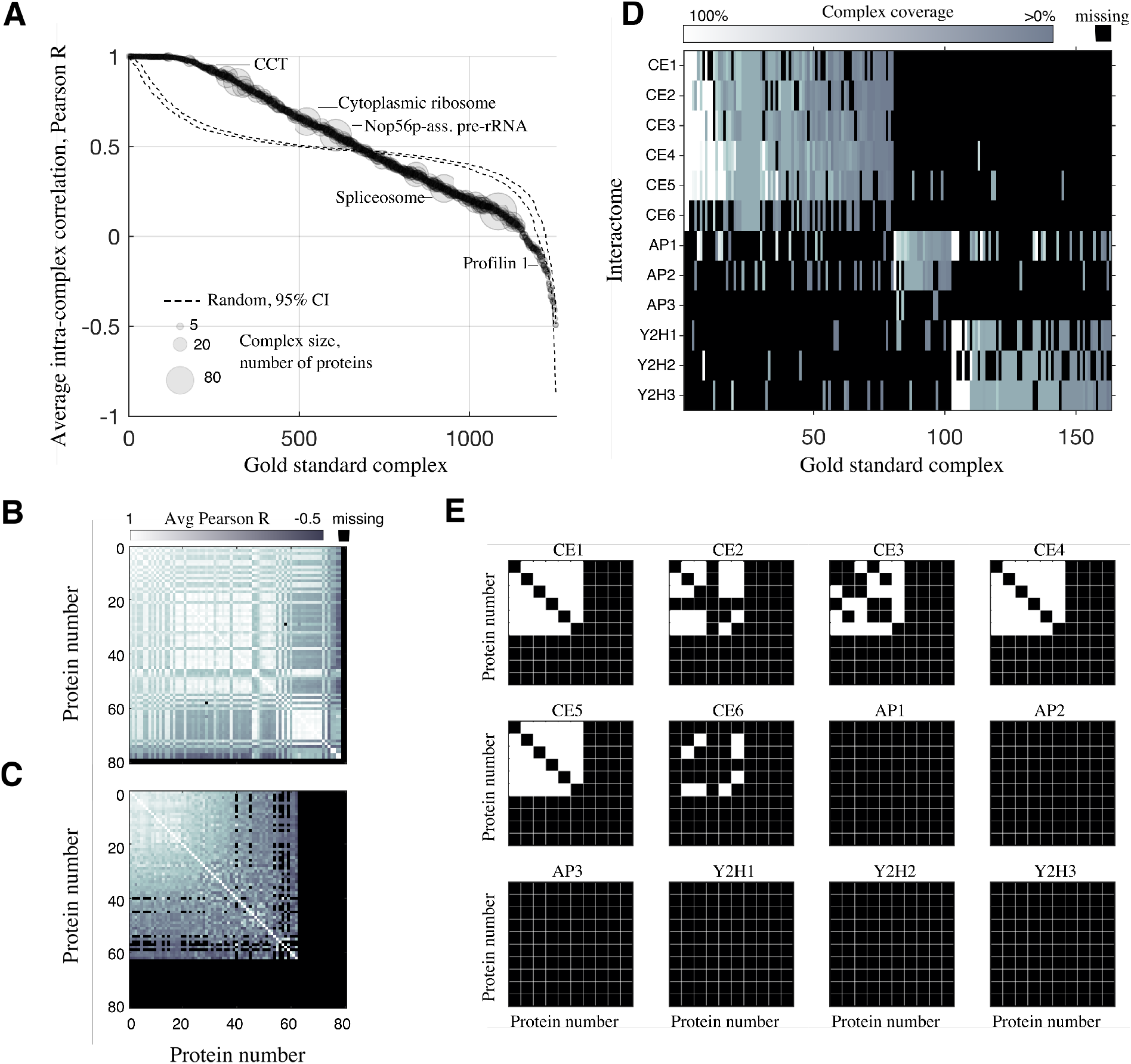
High-throughput techniques consistently recover some gold standard complexes and consistently fail to recover others. A. Average internal, pairwise correlation for every quantified gold standard complex. Only gold standard complexes with at least two identified proteins in one of 20 co-fractionation datasets are shown (1253/2652 CORUM complexes). Correlation values are pooled across the 20 co-fractionation datasets, and the number of internal, pairwise comparisons is given by marker size. The pattern expected by random chance is shown in red (95% CI). B. Connection matrix, cytoplasmic ribosome. Pairwise correlation values were averaged over 20 datasets. C. C complex spliceosome. D. Technique-specific gold standard complexes. All gold standard complexes predicted by at least 2/3 of the published interactomes from a given technique (CF, AP, Y2H), and no more than 1 interactome from the other techniques. E. Connection matrices for the Chaperonin Containing TCP-1 complex (CCT), a gold standard complex, taken from the published interactomes. White: interacting protein pair. Black: non-interacting.

Other high-throughput techniques display consistent over- and under-enrichment of specific gold standard complexes. Figure 3D shows gold standard complexes that were consistently predicted in one of three high-throughput techniques - CF, affinity purification mass spectrometry (AP-MS), or yeast two-hybrid (Y2H) - and largely absent from the others. Eighty gold standard complexes were predicted (≥1 interaction per complex) in 4/6 co-fractionation interactomes, while being predicted in no more than a single AP-MS or Y2H interactome (chance = 14 complexes, p = 0.005, bootstrap). Similarly, 61 gold standard complexes are predicted in at least 2/3 Y2H interactomes, while being predicted in at most one co-fractionation or AP-MS interactome (p < 0.001). Only 22 AP-MS-specific complexes are selected in this way (p = 0.49) possibly due to the low CORUM coverage of interactome AP3 (25). Technique-specific consistency is also seen at the level of pairwise interactions (Fig 3E).

In addition to being truly non-interacting, the absence of some gold standard complexes from published interactomes (Fig 3D) could result from low expression of interacting partners (rendering them difficult to quantify) or from none of the co-complex members being included as baits. To control for this, we additionally looked at the subset of gold standard complexes where at least one interaction could potentially be predicted in each study, as defined by having quantified proteins and baits (see Methods). The same pattern seen in Figure 3D persisted (Supp. Table 1), indicating that gold standard complexes seen by one method but not others cannot be explained by lack of coexpression or choice of bait, and therefore likely reflect the fact that the physical association of gold standard complexes is indeed conditionally dependent.

We note that the conditional dependence of gold standard complexes is not limited to the type of high-throughput experiment (CF, AP-MS, or Y2H). For example, using co-fractionation data where proteins were fractionated using a variety of techniques (2) (Supp. Table 1), the 60S ribosome gold standard complex consistently co-fractionated via sucrose fractionation (Supp. Fig 2A) but consistently failed to co-fractionate via heparin dual ion exchange (Supp. Fig 2B).

### 2.3 Universal gold standards improve interactome mapping

If a subset of database PPIs consistently fails to resemble interacting proteins for a given assay, performance should improve when these PPIs are removed from the gold standard set. We confirmed this was the case. We generated co-fractionation-specific gold standard subsets by selecting those complexes that were significantly enriched for interactions in interactomes CF4, CF5, and CF6 (2–4). Evaluating significance at four p-value thresholds (p < 1, 10^−2^, 10^−6^, 10^−10^) produced four subsets of CORUM complexes that contain 302, 122, 95, and 80 complexes, respectively (Table 1). To avoid training and testing on the same data, we defined the co-fractionation-specific gold standard subsets using interactomes published by other groups (CF4, CF5, CF6), and these gold standard subsets were then used to predict interactomes using co-fractionation data generated by our group.

These co-fractionation-specific CORUM subsets correspond significantly to housekeeping protein complexes. Using the Gini coefficient, a measure of inequality, we calculated consistency of mRNA expression (Fig 4A) (26) and protein expression (Fig 4B) (27) across tissues and cell types. As quantified by lower Gini coefficients, the expression levels of co-fractionation-specific complexes are significantly more consistent across tissues than other CORUM complexes (mRNA: Gini = 0.28 vs Gini = 0.40, p = 2.2 × 10^−16^, Welch two-sample t-test; protein: Gini = 0.40 vs Gini = 0.49, p = 2.2 × 10^−10^). This agrees with an orthogonal analysis of a mouse co-fractionation dataset collected by our group, which analyzed protein co-fractionation across seven tissue types. Quantifying co-fractionation via Pearson correlation, 15 CORUM complexes were found to be housekeeping complexes, as defined by average pairwise correlation significantly greater than chance in all seven tissues (p < 0.05, permutation test; Fig 4C). Of these 15 complexes, 8 overlap with the 122 co-fractionation- specific CORUM complexes, a significant overlap (p = 7.6 × 10^−8^, hypergeometric test; overlapping complexes marked by asterisk * in Figure 4C).

**Fig 4.**
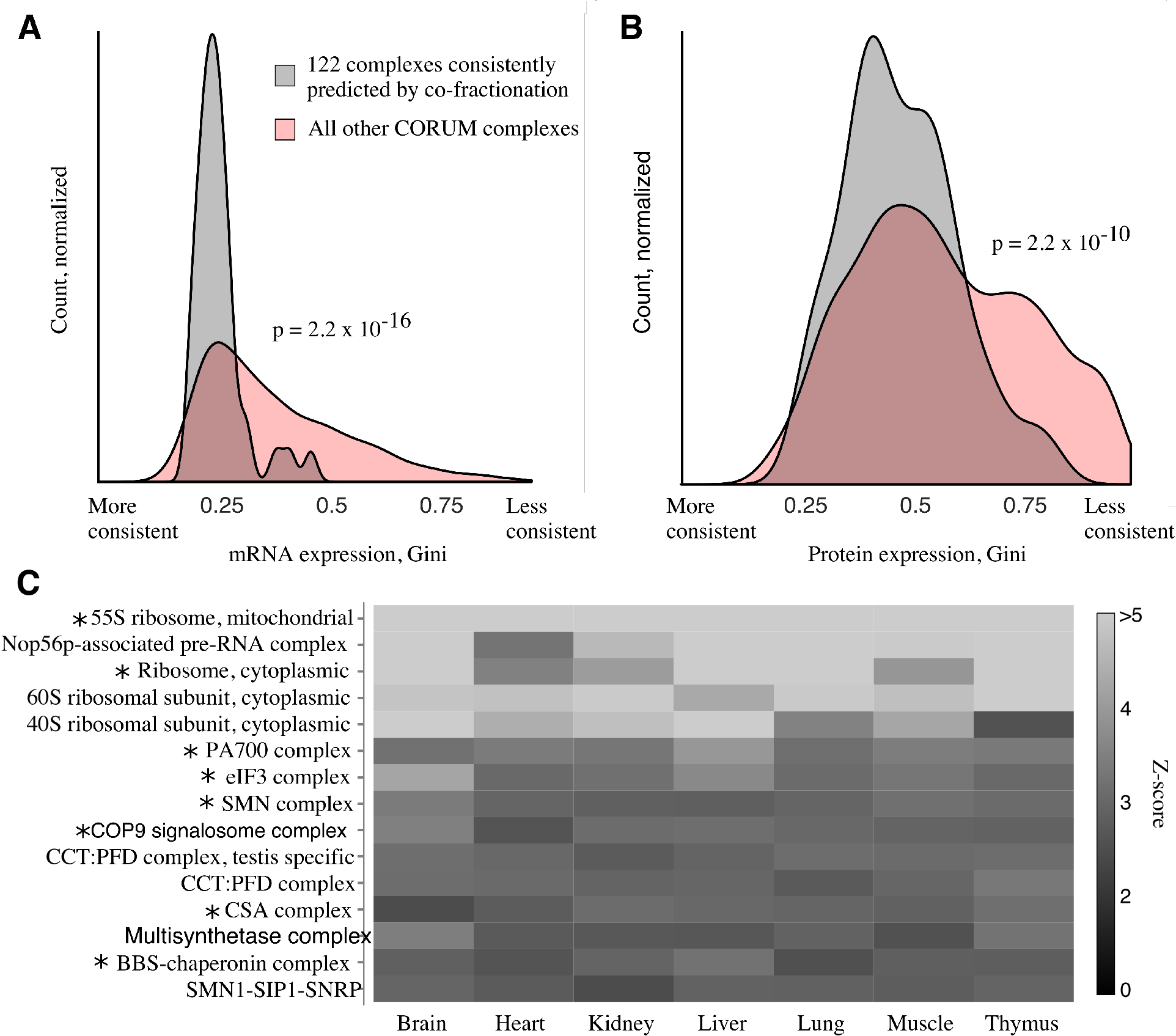
Gold standard complexes consistently predicted by co-fractionation correspond to housekeeping complexes. A. Consistency of mRNA expression levels across tissue types, Gini coefficient (26). B. Consistency of protein expression levels across tissue and cell types, Gini coefficient (27). C. Fifteen housekeeping CORUM complexes, defined by significant pairwise correlation between co-fractionation profiles in all seven tissues. The 8/15 complexes that overlap with the 122 complex subset of CORUM are marked by asterisks.

Using gold standard subsets generated in this way drastically alters the predicted interactomes (Fig 5). Controlling interactome quality via the ratio of true positives (TP) and false positives (FP), calculated as precision (TP/(TP + FP)), well-chosen gold standard subsets increased the size of the predicted interactome by up to an order of magnitude over randomly-chosen subsets (Fig 5A-C). Since FPs are defined as inter-complex protein pairs, they grow with the square of the gold standard set size. TPs, intra-complex pairs, grow linearly. Therefore there is a tendency for precision estimates to increase artificially as the gold standard set shrinks. For this reason we compared all cofractionation-specific subsets (Fig 5 black) to random subsets of the same size (red). Precision-recall curves, which visualize the tradeoff between quality and quantity of the interactomes, are also improved over random for increasingly stringent co-fractionation-specific gold standard subsets (Fig 5D-G).

**Fig 5.**
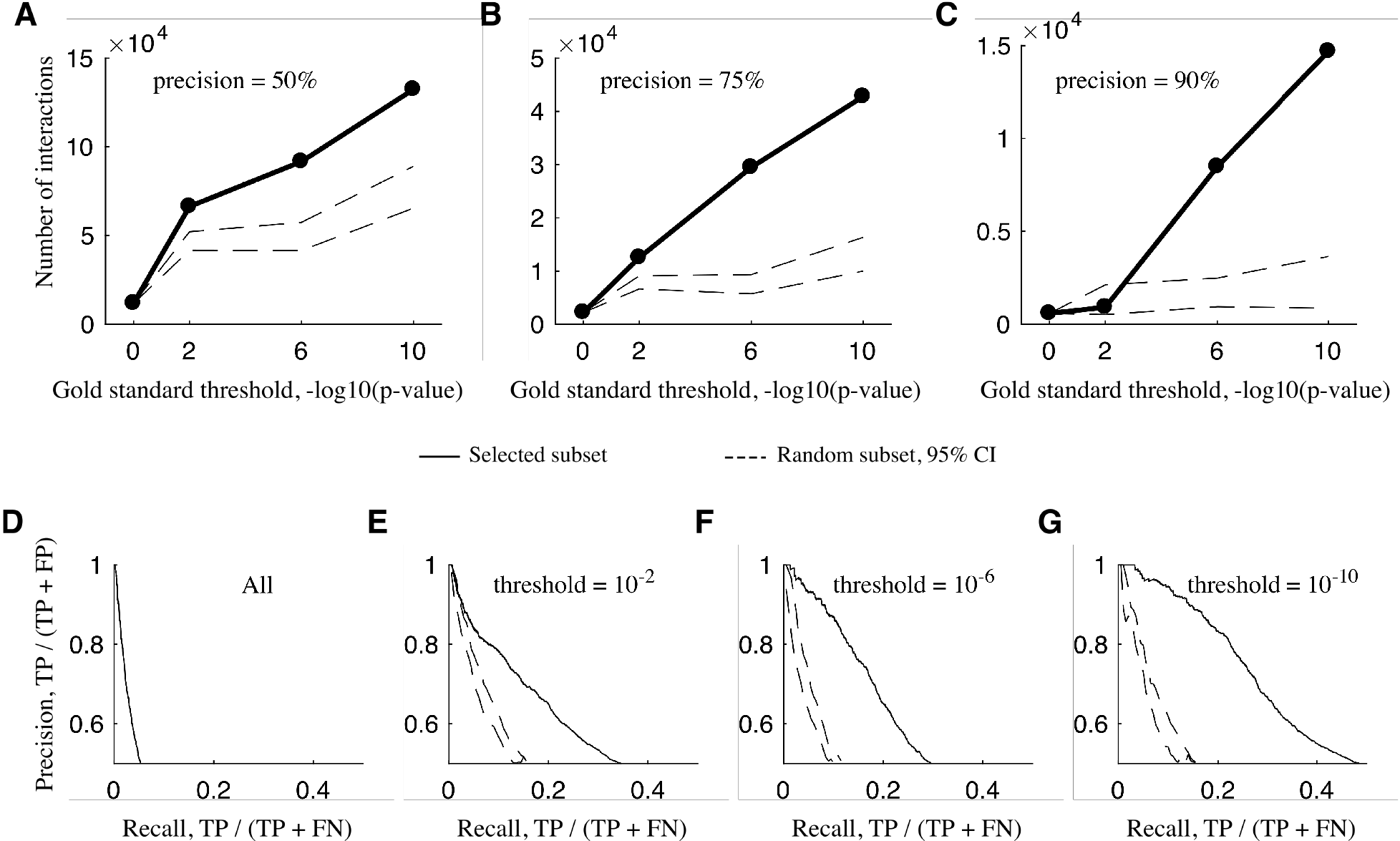
Using technique-specific gold standard subsets increases interactome size and/or quality. A. The size of interactomes produced by co-elution gold standard subsets of varying stringency (black) or randomly selected subsets of the same size as the co-elution specific subsets (red, 95% CI). Interactomes have 50% precision. B. 75% precision. C. 90% precision. D. Precision-recall curve for the interactome predicted using the entire gold standard set of interactions. E. Precision-recall curve predicted using the gold standard complexes that satisfied the 10^−2^ threshold. F. 10^−6^ threshold. G. 10^−10^ threshold. Precision-recall curves using random subsets of the same size are shown in red (95% CI).

## Discussion

Here we have estimated the discrepancies between interactome data generated by co-fractionation and curated gold standard interactions from the CORUM database. Across 20 datasets, 37% (54859/149477) of gold standard protein pairs display at most weak evidence for interaction (R < 0. 25, Pearson correlation), and 23% (34442/149477) show no evidence of interaction (R < 0) (Fig 1D, Fig 2). Other databases have a larger proportion of anti-correlated interactions, with up to 55% of database PPIs showing no evidence for interaction (Fig 2). Protein interaction networks have been compared elsewhere. For example, comparing the power of five PPI networks to predict cancer genes (28), benchmarking 21 networks for their ability to predict disease genes (29), and investigating their impact on recovering novel PPIs from high-throughput data (30). However, to our knowledge our study is the first to specifically address the conditional nature of PPI entries in these databases.

Since CORUM is manually curated from low-throughput experiments, we do not interpret these anticorrelated pairs as errors in the database. Rather, we attribute any discrepancy between our raw data and the databases to the conditional nature of protein interactions and the fact that databases compile evidence from many different experiments and conditions. Indeed, under certain conditions, 60S ribosomal proteins, which have been extensively studied and shown to interact, display poor evidence of interaction (Supp. Fig 1).

Therefore studies should take care not to conflate interaction databases, which attempt to list all interactions that *can* interact, with the subset of interactions that are in fact interacting in a given experiment. Doing so limits high-throughput interactome mapping studies. First, it artificially raises all estimates of error rates, since by definition a portion of the reference positive set is indistinguishable from the negative set. Second, when gold standard interactions are used as training labels in a classifier (2,3,7,11), classification accuracy will be reduced and fewer interactions and/or more noisy interactions will be predicted.

One solution is to use condition-or technique-specific gold standard subsets. We show that subsets of gold standard databases that have consistent, independent evidence taken from similar conditions to those under which the raw data was produced can increase the size of interactomes judged at the same precision level (Fig 4). We include this set of CORUM gold standard complexes and recommend it for future co-fractionation studies.

## Methods

### 4.1 Gold standards databases

We primarily used CORUM core complexes as an LC database of known protein complexes (Comprehensive Resource of Mammalian protein complexes, released February 2017) (31). CORUM is based entirely on experimentally verified interactions, all of which must have extensive low-throughput supporting data. To provide a broad sample of databases, we also analyzed interactions from an additional ten LC interaction databases: HPRD (release 9, last modified April 13, 2010) (16), MINT (downloaded June 8, 2017) (17), MENTHA (release June 5, 2017) (18), BIND (release 1.0, last modified May 20, 2014) (19), HIPPIE (release 2.0, last modified June 24, 2016) (20), IID (release April 2017) (21), BioGrid (release 3.4.149, accessed June 8, 2017) (32), PINA (version 2, last updated May 21, 2014) (22), HINT (version 4, downloaded June 8, 2017) (23), and DIP (release February 5, 2017) (24). We analyzed a subset of the full BioGrid database for which interactions were supported by at least two publications (N_full_ = 254886 interactions, N_subset_ = 39524). For databases such as CORUM that list complexes rather than pairwise interactions, gold standard interactions were defined as all protein pairs that are co-members of at least one gold standard complex. Only non-redundant, i.e. unique protein interactions were analyzed.

### 4.2 Co-fractionation profile datasets and mass spectrometry

The majority of co-fractionation data analyzed in this study was collected by our group and constitutes a broad sampling of SILAC-labelled co-fractionation datasets. Data was collected for four independent experiments, each mapping interactome rearrangements to an experimental treatment. Datasets in this study were separated by condition and replicate, such that an experiment with two conditions and three replicates would yield six datasets analyzed here. A total of 20 datasets are included in this study. Three experiments are previously published: two that map the reorganization of the HeLa interactome in response to stimulation with EGF (5) and *Salmonella enterica* infection (6), and one that maps the response of Jurkat T cells to Fas-mediated apoptosis (7). All fractionation was achieved by size exclusion chromatography except (7) which used blue-native page. Both methods separate protein complexes by molecular weight. The third co-fractionation dataset is available in this manuscript (Supp. Table 2). All co-fractionation data was quantified using SILAC ratios over successive fractions of a separation gradient, i.e. a chromatogram. Only protein chromatograms with quantification in five or more fractions were analyzed. In order to compare interactions seen by different fractionation techniques, we also analyzed previously published co-fractionation data generated by extensive biochemical fractionation (2). All co-fractionation profile datasets are composed of co-fractionation profiles, which are protein amount measured over successive fractions, quantified by mass spectrometry. There is one profile per protein or protein group for each combination of replicate and condition.

### 4.3 Published PPI interactomes

In addition to raw co-fractionation data, we analyzed twelve published human protein interactomes: three derived from co-fractionation data published by our lab (CF1 (7), CF2 (5), CF3 (6)), three derived from co-fractionation data not published by us (CF4 (3), CF5 (2), CF6 (4)), three AP-MS derived interactomes (AP1 (9), AP2 (10), AP3 (25)), and three Y2H interactomes (Y2H1 (12), Y2H2 (33), Y2H3 (34)). All interactomes were high-throughput and represent a broad sampling of the full human interactome.

### 4.4 Evaluating gold standard complexes

#### Raw co-fractionation profiles

To evaluate the degree to which gold standard PPIs are supported by co-fractionation data, we calculated the Pearson correlation coefficient and Euclidean distance between each pair of chromatograms in a dataset. For both measures, missing values in the chromatograms were replaced by zeros. When calculating Euclidean distance, all chromatograms were normalized to have a minimum value of 0 and a maximum value of 1. High correlation or low Euclidean distance was taken as evidence that the gold standard interaction was interacting in the sample.

#### Published interactomes

We mapped published pairwise protein-protein interactions to gold standard CORUM complexes. For each published interactome, we tested whether gold standard complexes were enriched for published interactions, meaning they contained significantly more pairwise interactions between complex members than the average rate (hypergeometric test). We took significant enrichment as evidence that the published interactome supported the gold standard complex. To standardize interactomes with each other and CORUM, all isoform tags were removed from protein IDs.

To control for expression and different baits, we also defined the subset of gold standard complexes in each study in which at least one interaction could be predicted. For CF interactomes this was defined as a complex in which at least two co-complex members are present in the raw data (raw data for (3) downloaded here http://metazoa.med.utoronto.ca/; (4) and (2) raw data taken from publication). For AP-MS interactomes we assumed the matrix model, meaning that a bait protein need not be present in a gold standard complex for an interaction to be predicted in the gold standard complex, as long as a gold standard complex member is associated with a bait protein. Therefore if at least two members of the gold standard complex were present in the AP-MS interactome, we defined that gold standard complex as a complex that could be predicted by the study. Finally, for Y2H interactomes we defined a gold standard complex as able to be predicted if at least one complex member was a bait protein.

### 4.5 Co-fractionation-specific gold standard subsets

In this study, we used subsets of the gold standard complexes that are consistently supported by cofractionation interactomes. For these co-fractionation subsets, we used all CORUM complexes that were significantly enriched for interactions (hypergeometric test) in CF4, CF5, and CF6. Significance was assessed at a range of p-value thresholds: 1, 10^−2^, 10^−6^ and 10^−10^. A threshold of p = 1 produced the subset of CORUM complexes with at least one interaction in CF4, CF5, and CF6. The number of CORUM complexes (interactions) in each subset were 302 (33378), 122 (10953), 95 (6326) and 80 (4861), respectively.

To control for the effects of simply reducing the size of the gold standards, we generated random subsets of gold standard PPIs with the same size as the selected subsets. Of the full 46413 unique PPIs in the core CORUM complexes, we randomly sampled 33378, 10953, 6326, and 4861 PPIs without replacement. Random sampling was repeated 100 times for each of the subset sizes, and interactomes were predicted using each random subset with the PrInCE software package.

### 4.6 Interactome prediction

For this study, interactomes were predicted using the PrInCE software (Predicting Interactomes via Co-Elution), a software package developed by our lab for the analysis of co-fractionation datasets (11). PrInCE measures the similarity between every pair of co-fractionation profiles using a variety of similarity measures, such as Pearson correlation and Euclidean distance. Gold standard interactions are used as true positive labels (TP) in a Naive Bayes classifier. False positive interactions (FP) are defined as interactions between a pair of proteins that both occur in the gold standard database, but are not members of the same gold standard complex, e.g. an interaction between a ribosomal protein and a proteasomal protein. PrInCE assesses the quality of the predicted interactome using precision, where precision = TP/(TP + FP).

## Acknowledgements

We thank N. Brown, C. Kerr, A. McAfee, and A. Nastasa for critical suggestions. M.A.S. is supported by a CIHR Frederick Banting and Charles Best Canada Graduate Scholarship, a UBC Four Year Fellowship, and a Vancouver Coastal Health-CIHR-UBC MD/PhD Studentship Award.

## Supporting information

**S1 Fig. Restricting gold standard PPIs to those supported by two or more publications does not eliminate uncorrelated protein pairs, as measured by Pearson correlation R<0**. Each point is one dataset. Horizontal lines show medians. Red: all non-gold standard protein pairs. Black: non-redundant gold standard pairs. “All pairs” and “BioGrid” correspond to Figure 1.

**S2 Fig. 60S ribosome co-fractionates via sucrose fractionation (A) but not via heparin dual ion exchange (B)**. Pearson R. Plots show replicates. Missing (black) are protein pairs where neither protein was detected.

**S1 Table. Some CORUM complexes are predicted by a single high-throughput technique, as measured by average complex coverage**. Complex coverage = number of pairwise interactions in a published interactome / total pairwise connections within a complex. Parentheses show number of complexes averaged. CF-specific complexes correspond to numbers 1-80 in Figure 3D, AP/MS-specific to 81-102, and Y2H-specific to 103-163. To control for expression and bait selection, only complexes that could be be predicted in each interactome are included (see Methods).

**S2 Table. Third co-fractionation dataset.**

